# A whole-brain and cross-diagnostic perspective on functional brain network dysfunction

**DOI:** 10.1101/326728

**Authors:** Marjolein Spronk, Kaustubh Kulkarni, Jie Lisa Ji, Brian P. Keane, Alan Anticevic, Michael W Cole

## Abstract

A wide variety of mental disorders have been associated with resting-state functional network alterations, which are thought to contribute to the cognitive changes underlying mental illness. These observations have seemed to support various theories postulating large-scale disruptions of brain systems in mental illness. However, existing approaches isolate differences in network organization without putting those differences in broad, whole-brain perspective. Using a graph distance measure – connectome-wide correlation – we found that whole-brain resting-state functional network organization in humans is highly similar across a variety of mental diseases and healthy controls. This similarity was observed across autism spectrum disorder, attention-deficit hyperactivity disorder, and schizophrenia. Nonetheless, subtle differences in network graph distance were predictive of diagnosis, suggesting that while functional connectomes differ little across health and disease those differences are informative. Such small network alterations may reflect the fact that most psychiatric patients maintain overall cognitive abilities similar to those of healthy individuals (relative to, e.g., the most severe schizophrenia cases), such that whole-brain functional network organization is expected to differ only subtly even for mental diseases with devastating effects on everyday life. These results suggest a need to reevaluate neurocognitive theories of mental illness, with a role for subtle functional brain network changes in the production of an array of mental diseases.

## Significance Statement

The devastating effects of mental diseases on everyday life can lead one to assume that substantial disruptions to brain organization underlie such diseases. Indeed, many theories postulate large-scale brain network disruptions driven by widespread neurotransmitter dysfunctions. Yet most patients maintain most basic cognitive abilities (broadly construed to include perceptual, attentional, memory-based, emotional, and motor capabilities) enjoyed by healthy individuals. We therefore hypothesized (and confirmed) that overall brain network organization – thought to be essential in enabling cognitive abilities – is highly similar between patients and healthy individuals across a variety of mental disorders. These results provide hope that small, well-targeted alterations to brain network organization may provide meaningful improvements for a variety of mental disorders.

## 1. Introduction

Cognitive dysfunction (broadly construed to include perceptual, attentional, memory-based, emotional, and motor capabilities) is seen in a range of mental disorders and can significantly impact a patient’s well-being. As a neural correlate of cognitive impairment (Greicius, 2008; Zhang and Raichle, 2010), abnormal resting-state functional connectivity (RSFC) has been used to identify neural mechanisms underlying mental illness (for review see Cole et al., 2014b). RSFC has strong potential for providing important insights in this area, given its relationship with a variety of cognitive abilities (Cole et al., 2011; Smith et al., 2015; Shen et al., 2017), its generalization to a variety of task states (Cole et al., 2014a; Krienen et al., 2014), and recent findings indicating RSFC describes the routes of cognitive information flow during task performance (Cole et al., 2016; Ito et al., 2017). Indeed, this method has already provided promising results in the search for biomarkers in several psychiatric disorders. For example, combined with graph theory measures, machine learning approaches have shown the ability to classify patient and control subjects successfully for a range of disorders (Yahata et al., 2017).

Many theories postulate large-scale disruption of brain systems in psychiatric disorders (Uhlhaas and Singer, 2011), typically driven by widespread neurotransmitter dysfunction (Olney et al., 1999; Risch et al., 2009; Nakic et al., 2010; M. et al., 2013). However, such large-scale disruptions of brain systems have not been thoroughly tested. This is largely due to a fundamental aspect of methodology used in clinical studies, wherein only differences among patients and healthy controls are emphasized rather than both differences and similarities. Yet taking similarities into account is essential for gaining perspective on the nature and severity of the neural changes underlying mental disorders. For instance, it is possible that a widespread neurotransmitter dysfunction causes a given disease, but only by subtly altering overall network organization. This would be important to know for understanding the underlying causes of the disease and for developing treatments.

Here, we therefore take a comprehensive whole-connectome perspective using a simple graph distance measure – connectome-wide correlation (Schultz and Cole, 2016) – that quantifies changes in all functional connections at once, providing information about the overall pattern of intrinsic network architecture. In this way, we were able to compare RSFC patterns across several psychiatric disorders and healthy control groups and derive measures of functional network pattern (dis)similarity. We initially developed this approach to compare task-evoked functional connectomes to resting-state functional connectomes (Cole et al., 2014a; Schultz and Cole, 2016), but we apply it here to compare clinical connectomes to healthy control connectomes in resting state. This provides a new way of quantifying brain function and neural system organization in psychiatric disorders.

We (and others) recently found that task-related changes to functional network organization are small relative to the overall functional network organization during rest and a variety of tasks (Cole et al., 2014a; Krienen et al., 2014). This suggests that meaningful changes in cognition (in this case task-related cognitive differences) correspond with small functional network changes. Further, the majority of patients appear to maintain most basic cognitive abilities enjoyed by healthy individuals (e.g., the ability to recognize common objects, navigate a room, produce speech, read simple sentences). Building on this logic, we hypothesized that the overall intrinsic functional network architecture – which appears to support these cognitive abilities via its particular network organization (Smith et al., 2009; Cole et al., 2016; Ito et al., 2017) – would be very similar across patients, and between patient and control subjects. We tested this idea across a range of patient datasets varying in diagnosis, age and symptom severity. Importantly, even with high cross-group similarity, between-group differences can still indicate meaningful changes in functional network architecture. To test this, we predicted group membership (clinical vs. control) for each participant using a classification analysis based on RSFC pattern similarity to either group.

Testing these hypotheses about whole-brain intrinsic architecture pattern similarity across disorders – here including autism spectrum disorder (ASD), attention-deficit hyperactivity disorder (ADHD), and schizophrenia – would enable us to put disease-related RSFC alterations in perspective, and possibly help shape future theories of the neural basis of cognitive deficits in these populations.

## 2. Methods

### 2.1 Datasets and participants

Subjects from four different datasets were included in the present study to facilitate generalization of findings across mental disorders. ADHD subjects are part of the publicly available ADHD-200 dataset from NYU Langone Medical Center (http://fcon1000.projects.nitrc.org/indi/adhd200/), which includes 123 patients diagnosed with ADHD and 99 control subjects (presence or absence of an ADHD diagnosis based on evaluations with the K-SADS-PL (Kaufman et al., 1997) and the CPRS-LV (Gurley, 2011)). Subjects with low-quality resting-state or anatomical data, as determined by the ADHD-200 initiative’s quality assessment based on visual time series inspection, were excluded from our sample. Resting-state and anatomical data of the remaining 87 healthy control subjects and 93 ADHD subjects were preprocessed (age range 7-18 yrs). ASD subjects were part of the publicly available ABIDE dataset from NYU Langone Medical Center (http://fcon1000.projects.nitrc.org/indi/abide/abideI.html). Inclusion as patient in the study required a clinician’s DSM-IV-TR diagnosis of Autistic Disorder, Asperger’s Disorder, or Pervasive Developmental Disorder Not-Otherwise-Specified, which was supported by review of available records, an Autism Diagnostic Observation Schedule, review of the participant’s history, and when possible, an Autism Diagnostic Interview-Revised. Following these inclusion criteria, 79 ASD and 105 control subjects were included for preprocessing (age range 6.5-39.1 years). The ADHD and ASD samples in this study were collected at the same site, and could therefore be directly compared in our analyses without site differences as confounds.

In addition to these two samples with neurodevelopmental disorders, two schizophrenia samples were included in the present study. The first was collected at Yale University and consisted of 90 schizophrenia patients and 90 healthy control subjects (age range 17-65). Patients were identified through outpatient clinics and community mental health facilities (Anticevic et al., 2013). The second sample is the publicly available COBRE schizophrenia dataset, with 72 schizophrenia patients and 75 healthy control subjects (age range 18-65). Each patient completed the Structured Clinical Interview for DSM-IV Axis I disorders (First et al., 2002) to confirm their diagnosis (see http://fcon1000.projects.nitrc.org/indi/retro/cobre.html for exclusion criteria).

The number of subjects in the final sample are displayed in Table 1. Motion scrubbing/censoring was applied (see Section 2.3 Preprocessing), given that it is thought to be especially important for RSFC analyses of clinical groups with high levels of head motion (Power et al., 2014). We scrubbed time points above a standard framewise displacement (FD) threshold of 0.5 mm. We additionally excluded subjects with less than 4 minutes of resting-state fMRI data remaining after motion scrubbing to increase the likely reliability of the functional connectivity estimates in our sample. This resulted in 10 control subjects and 12 patients being removed for analysis from the ADHD dataset, 3 control subjects and 9 patients from the ASD dataset, 6 control subjects and 15 schizophrenia subjects from the Yale dataset, and 29 control subjects and 38 schizophrenia subjects from the COBRE dataset. Hence, subject retention rates after scrubbing for control subjects and patients were respectively 89% and 87% for ADHD, 97% and 98% for ASD, 93% and 83% for the Yale schizophrenia data, and 60% and 46% for the COBRE schizophrenia data.

The data were collected by multiple research groups and the type of clinical information collected was dependent on diagnosis (ADHD vs ASD vs schizophrenia). Consequently, diagnosis-specific symptom scores were available for each dataset: ADHD-index for ADHD patients, Autism Diagnostic Observation Schedule (ADOS) for ASD, and Positive and Negative Symptom Scale (PANSS) for Schizophrenia were included (see Table 1 for mean symptom scores). As expected, the mean ADHD index is higher in ADHD patients than control subjects (t(1,153)=−20.75, p<0.0001, d=3.4). No symptom scores were available for the other control groups. Comorbidity was reported for 25 ADHD patients and 26 ASD patients, and no comorbidity data were available for the schizophrenia groups.

Since our main focus in this study is on similarity of FC architecture across patients and healthy control subjects regardless of variation in phenotypic factors, groups were not specifically matched on age, gender and IQ, although matching for age and gender was done at time of collection for all available datasets. In the final sample, lower IQ scores were found in ADHD (t(1,146)= 1.99, p=0.05, d=0.33), ASD (t(1,170)=2.65, p=0.01, d=0.34) and Yale schizophrenia patients (t(1,148)=4.28, p<.001, d=0.73) than in control subjects of those groups. No IQ scores were available for the COBRE schizophrenia sample. However, education scores were not significantly different between patients and control subjects (t(1,72)=1.74, p=0.09, d=0.37)). ADHD subjects in the final sample for analysis were slightly younger than controls (t(1,148)=2.2, p=0.03, d=0.34), but there were no significant age differences in the other data samples.

**Table 1.**
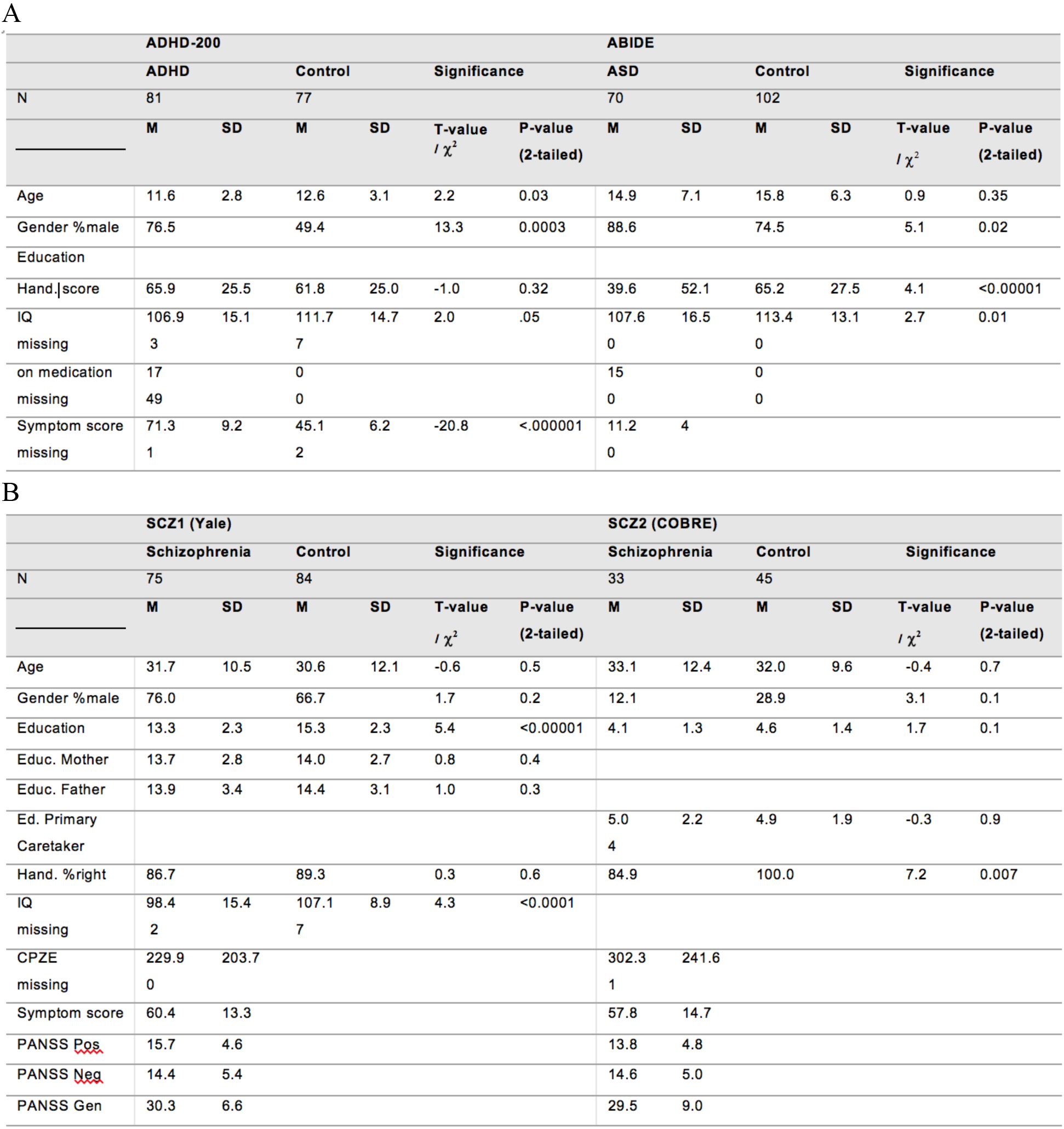
Clinical and demographic characteristics

### 2.2 Neuroimaging acquisition

The resting-state fMRI data for ADHD and ASD patients and healthy control subjects were acquired on a Siemens Magnetom Allegra 3.0 Tesla MRI scanner. Participants were asked to remain still with their eyes closed, but without falling asleep. For the resting state multi-echo EPI images, 33 slices were acquired every 2,000 ms (FOV = 240 × 192 mm, TE=15 ms, flip angle = 90°, voxel size 3 × 3 × 4 mm^3^) with a total of 180 volumes per run. Functional images for the Yale dataset were collected using a 3-T Siemens Allegra scanner. 210 volumes were acquired for each participant in the Yale schizophrenia dataset, with a TR of 1500 ms, fov= 220 mm, TE = 27 ms, flip angle = 60°, and voxel size 3.43 × 3.43 × 4, eyes open (Anticevic et al., 2014). For the COBRE dataset, a 3-T Siemens Trio Tim scanner was used to acquire 33 slices every 2,000 ms (TE = 29ms, flip angle = 75°, fov = 240 mm, voxel size 3 × 3 × 4 mm^3^) with a total of 150 volumes per run. Participants were asked to keep their eyes open. A high resolution T1-weighted MPRAGE was collected for each participant.

### 2.3 Preprocessing

Preprocessing was performed using Freesurfer (to identify ventricles, white matter, gray matter and anatomical structures) (Destrieux et al., 2010), FSL’s FLIRT for brain image alignment (Smith et al., 2004), and AFNI (Cox, 1996) for all other preprocessing steps. Volume analysis was performed on all datasets. Functional images underwent slice-time correction, alignment (with the anatomical image and MNI template), removal of first 6 s of data, motion scrubbing (censoring volumes with high motion based on framewise displacement with a 0.5 mm threshold, see Power et al., 2012), time series extraction, nuisance regression (removal of 6 motion estimates, ventricle and white matter signals, and their derivatives), bandpass filtering (0.008-0.09Hz), and spatial smoothing (FWHM = 6mm).

An additional cortical surface-based analysis was performed with the Yale schizophrenia sample (which was also preprocessed using the volume-based approach). The surface-based analysis involved using the HCP minimal preprocessing pipeline (Glasser et al., 2013), followed by removal of the first 6 seconds of data, motion scrubbing (using framewise displacement > 0.5 mm), time series extraction, nuisance regression, and temporal filtering. Note that obtaining similar results to the main analyses with this distinct preprocessing stream would indicate robustness of results to particular analysis choices.

### 2.4 Statistical Analysis

For our main analysis including all datasets, we sampled data for each individual from a set of 264 independently-identified functional regions (Power et al., 2011) to provide results at the region, system, and whole-brain level. In an additional analysis – including only schizophrenia patients from the Yale study – we used another set of regions with 360 parcels (Glasser et al.,2016) to test the robustness of our results, see section 3.4. For both analyses, time courses were extracted from each region and averaged across voxels/vertices for use in all subsequent analyses, which were performed in MATLAB 2014b (The Mathworks). Pearson correlations were calculated between all ROIs for each subject, and the resulting functional connectivity (FC) matrix was Fisher’s Z-transformed to normalize the distribution of values. These values were then used on all subsequent statistical tests.

To determine similarity of whole-brain RSFC between two groups, FC patterns were compared by taking the upper triangle of the RSFC matrices to be compared (thereby excluding self-connections and redundant connections), vectorizing the Fisher’s Z-transformed FC values, and computing Spearman rank correlations (and, separately, Pearson correlations) on the resulting vectors. Note that Pearson correlation is a standard pattern distance approach, with a simple transform (1-r) switching the measure from similarity to dissimilarity. We chose to use this distance measure in terms of similarity rather than dissimilarity (despite their one-to-one correspondence) in order to facilitate intuitive understanding of the results. Further, using the original r values (and rho values in the case of Spearman correlations) allowed us to use Fisher’s z-transform in order to make better statistical inferences at the group level.

Since region-level differences have been found in RSFC between the clinical populations studied here and healthy controls, we also compared individual connections between groups. T-tests were run on the group averages of each pair of regions (i.e., 34,716 connections). Due to the large number of statistical tests, we report false discovery rate (FDR)-corrected p-values (Genovese et al., 2002).

We also ran a classification analysis (Yahata et al., 2017) to further investigate if existing differences in resting-state network architecture can be used to distinguish patients from healthy controls. In order to investigate whether we could accurately predict group membership based on RSFC patterns, we conducted a leave-two-out classification analysis for each dataset separately to classify each subject. We balanced datasets by randomly selecting the same number of patients and control subjects per dataset (based on the number of subjects in the smallest group). Such balancing is essential to reduce classifier bias (e.g., always predicting “patient” for a dataset with more patients than controls). For each cross-validation fold, we compared one patient and one healthy control subjects’ RSFC patterns (test set) to the average patient and average control RSFC pattern (training set, excluding the held-out control and patient). In addition, we tested across ADHD and ASD, and across schizophrenia datasets by training on one dataset and testing on the other dataset, and vice versa. Similarity of a subject’s RSFC pattern to the average RSFC pattern of the other dataset’s groups was used to assign a label “patient” or “control” to a subject in the test set. Prediction accuracy, sensitivity and specificity – based on a classification with 10,000 randomly-initialized iterations – are reported in the results section.

## 3. Results

### 3.1 Large overlap in RSFC patterns in ADHD, ASD patients and healthy controls

We used fMRI to examine RSFC in patients with various mental disorders. We sought to identify common functional network patterns (compared to matched healthy control subjects) for multiple mental disorders with different symptomatology. We started by analyzing publicly-available resting-state fMRI data from ADHD and ASD patients, and healthy control subjects. We extracted time series from 264 regions of interest (Power et al., 2011, see Fig. 1A) and calculated all pairwise correlations, which were normalized with a Fisher’s Z transformation for all subsequent statistical tests. RSFC matrices in network order for ADHD and ASD are displayed in Fig. 1B (we discuss the results with schizophrenia in the following section).

**Fig. 1.**
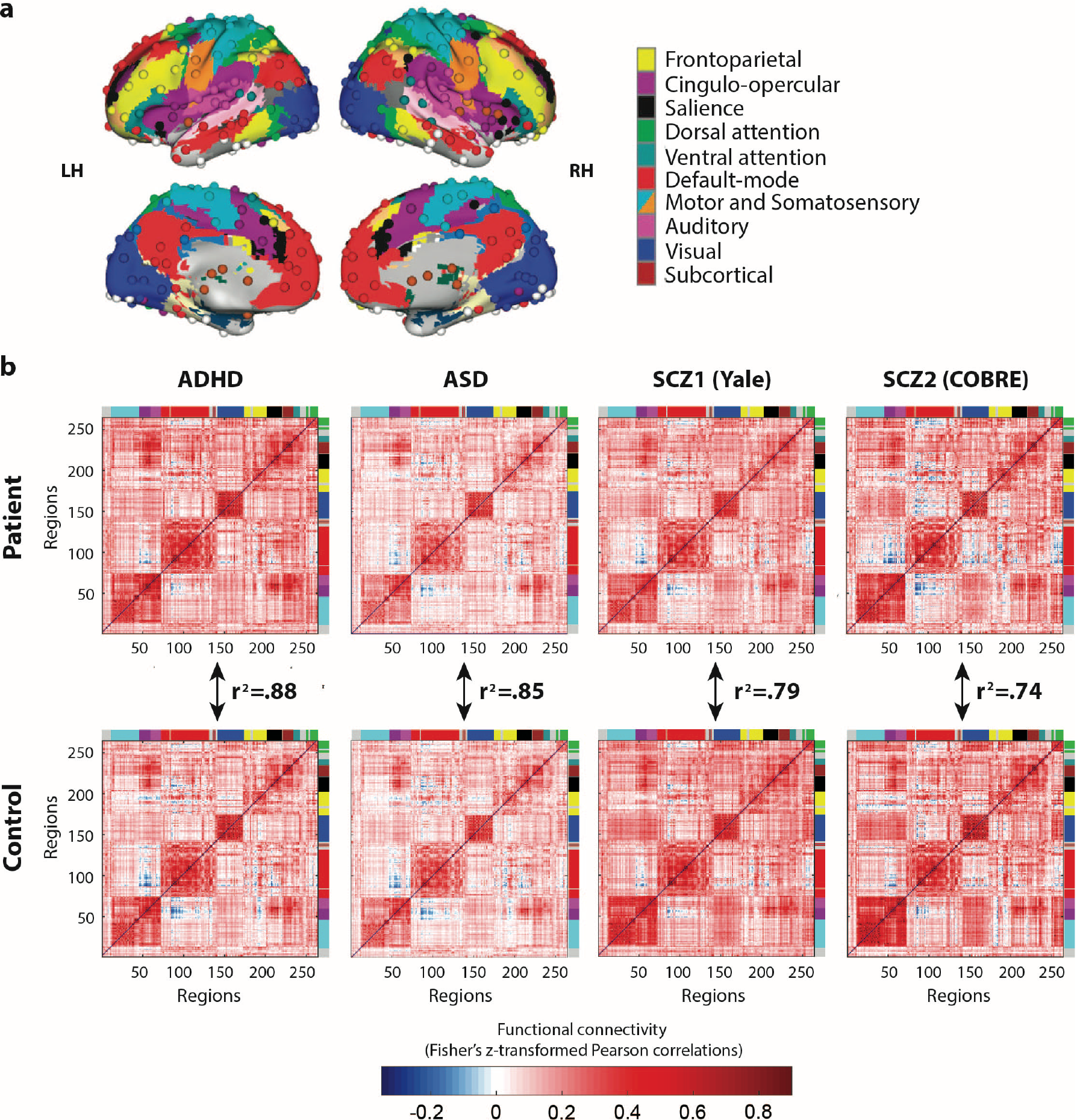
Whole-brain functional network organization is highly similar between patient and healthy control groups across a variety of mental diseases. **A)** Functionally-defined cortical and subcortical regions (nodes), along with a standard network partition based on healthy young adults (Power et al., 2011) used for whole-brain analyses. **B)** RSFC matrices for ADHD, ASD and schizophrenia patients show highly similar resting-state network architectures with healthy individuals. Colors along the edges of the matrices indicate previously-identified functional networks, matching the colors in panel A. The red blocks along the diagonal indicate stronger RSFC within relative to between networks. The R^2^ values can be interpreted as the percentage of linear variance shared (e.g., 0.88 = 88% shared linear variance). Note that these R^2^ values can easily be converted into correlation distance (1-r) values, such that these results also imply low distance in network state space (Schultz and Cole, 2016) between groups.

For both clinical studies, we calculated similarity of RSFC patterns between patients and control subjects using the 264×264 RSFC matrices with pairwise connections. Each individual’s FC matrix consisted of a minimum of 4 minutes of resting-state fMRI data (after removing high-motion volumes; see Methods). A non-parametric Spearman rank correlation (rho) for the whole-brain RSFC configuration was used to compare patients’ RSFC pattern to that of control subjects.

We found that for both clinical groups RSFC patterns were highly similar to the control RSFC patterns. ADHD & control: Spearman rho = 0.94, *p* < 0.00001; Autism Spectrum Disorder & control: Spearman rho = 0.92, *p* < 0.00001. These results suggest a largely common intrinsic network architecture in healthy individuals and patients. The R^2^ was .88 for ADHD and .85 for ASD, suggesting 88% and 85% of the linear variance was shared between patients and controls on average for these groups. Results were similar when using Pearson correlations rather than Spearman correlations. ADHD & control: Pearson r= 0.96, *p* < 0.00001; Autism Spectrum Disorder & control: Pearson r= 0.95, *p* < 0.00001. By focusing on the whole-brain RSFC pattern – using a connectome-wide comparison measure – we found that intrinsic network architecture is highly similar across patients and healthy control subjects in these two clinical groups, as hypothesized.

### 3.2 Large overlap in RSFC patterns in schizophrenia and healthy controls

Since the ADHD-200 NYU and ABIDE NYU data were collected at the same site, in the same age group, and with the same MRI parameters, we decided to test the generalizability of our conclusions with a distinct mental disorder and distinct datasets. These datasets included schizophrenia patients and matched healthy controls collected at two separate sites, in a distinct age group, and with distinct MRI parameters from the previous datasets. Whereas comorbidity exists between schizophrenia and ASD, and between schizophrenia and ADHD, there is much less overlap in symptoms than between ASD and ADHD (note that a dual diagnosis of these disorders is only possible since the introduction of DSM-V). Moreover, in this group of patients and control subjects only adults were included, in contrast to the ADHD (only children) and ASD (mostly children, but also adults) groups.

Our connectome-wide distance analysis again indicated a highly similar RSFC configuration pattern between schizophrenia patients and healthy control subjects with R^2^ of 0.79 and 0.74 (respectively, Yale schizophrenia & control: Spearman rho = 0.89, *p* < 0.00001; COBRE schizophrenia & control: Spearman rho = 0.86, *p* < 0.00001, see Fig. 1B), demonstrating that the previous finding of RSFC similarity generalizes to other mental disorders (and other age groups and MRI parameters). Results were similar when Pearson correlation was used as the similarity/distance measure (Yale schizophrenia & control: r = 0.92, *p* < 0.00001; COBRE schizophrenia & control: r = 0.90, *p* < 0.00001).

A combined analysis of the two schizophrenia datasets resulted in even higher similarity between patient and control groups with an R^2^ of .83 (schizophrenia & control: Spearman rho = 0.91, *p* < 0.00001; Pearson r = 0.94, *p* < 0.00001).

### 3.3 RSFC pattern similarity across disorders

We next sought to determine the general similarity between patients and healthy controls across all three mental disorders. The whole-brain RSFC matrices were averaged across all patients and, separately, across all healthy controls. These matrices showed a strong correlation between patients and controls (Spearman rho = 0.96, *p* < 0.00001; Pearson r = 0.97, *p* < 0.00001). The R^2^ was 0.92, suggesting 92% of the linear variance was shared between patients and controls on average. Similarity between patient groups (ADHD/ASD: rho = 0.92, *p* < 0.00001; ADHD/schizophrenia: rho = 0.83, *p* < 0.00001; ASD/schizophrenia: rho = 0.83, *p* < 0.00001) and between control groups (ADHD/ASD: rho = 0.95,*p* < 0.00001; ADHD/schizophrenia: rho = 0.84, *p* < 0.00001; ASD/schizophrenia: rho = 0.86, *p* < 0.00001) was high as well.

### 3.4 Replication of architecture similarity with a different set of ROIs

To test the robustness of our RSFC similarity findings, we sought to replicate them with a different set of brain regions (with the Yale schizophrenia dataset). We used a parcellation of functionally-defined regions that was recently developed by Glasser et al. (2016). This cortical parcellation with 360 regions was constructed using convergence across multiple neuroimaging techniques (resting-state and task fMRI, myelin maps and cortical thickness), and is believed to be more accurate than previous parcellations because of the consistency of areal borders between data from different imaging modalities (Glasser et al., 2016). Networks were defined in a separate study using community detection analysis with RSFC data (see Methods section 2.4) (Spronk et al., 2017). Similar to the analysis with volume (Power) regions, data from surface regions were extracted, RSFC estimated, and Spearman correlations between RSFC matrices were calculated. Similarity analysis of RSFC matrices of schizophrenia patients and control subjects returned R^2^ = 0.85 for the Yale dataset (rho = 0.92, *p* < 0.00001, see Fig. 2; RSFC similarity was R^2^ = 0.79 with the volume ROIs), again supporting the existence of a largely similar intrinsic functional network architecture across patients and controls.

**Fig. 2.**
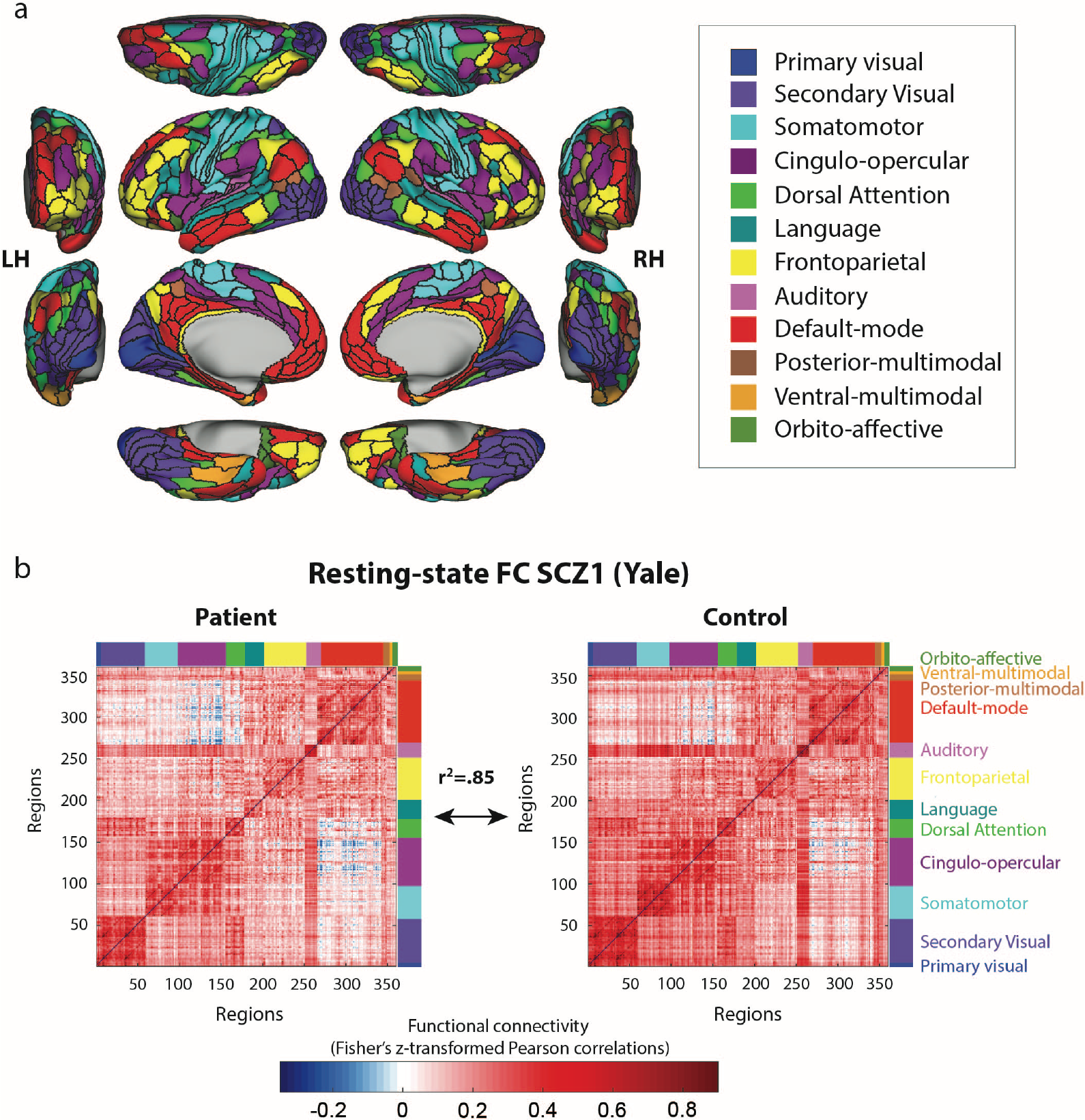
Testing for generalization to a distinct set of regions, network definition. **A)** Network partition based on the multi-modal parcellation by Glasser et al. (2016). The main parcellation (Fig. 1) used regions defined in terms of voxels, whereas this parcellation is defined in terms of surface vertices. Functional networks were assigned based on the General Louvain method for community detection with resting-state data in healthy adults (Spronk et al., 2017). **B)** RSFC matrices based on this alternative set of regions and networks were used to replicate the previously-found similarity between schizophrenia patients and healthy control subjects. This suggests the particular choice of regions and networks used for the main analyses did not substantially influence results – that the results are generalizable.

### 3.5 Small-scale RSFC deviation in mental disorders

Importantly, similar RSFC patterns between patient and control groups doesn’t necessarily mean that all functional connections are normal in patients with mental disorders. Indeed, many results in the literature demonstrate statistically significant RSFC differences between patients and healthy controls, even after controlling for between-group motion confounds as we have here. To test for consistency with prior studies, we also compared individual connections, i.e., tested for between-group differences on every unique connection pair in the matrix (34,716 connections). In this way, in addition to testing the overall pattern with Spearman correlations, we examined smaller-scale (individual connection-level) FC alterations in the clinical groups. The resulting patterns of t-values (patient vs. control for each connection) are displayed in Figure 3a. These patterns of t-values showed similarity across the datasets with a significant correlation between t-value patterns of ADHD and ASD datasets (rho = 0.25, *p* < 0.000001) and the two schizophrenia datasets (rho = 0.29, *p* < 0.000001). Between ADHD/ASD and schizophrenia datasets, correlations were weaker but also significant (ADHD & Yale: rho = -0.04, *p* < 0.000001, ASD & Yale: rho = 0.14, *p* < 0.000001, ADHD & COBRE: rho = 0.04, *p* < 0.000001, ASD & COBRE: rho = 0.09, *p* < 0.000001). After thresholding (*p* < 0.05) and FDR correcting the t-value patterns for multiple comparisons, we found several connections that had significantly altered RSFC connectivity in ADHD and schizophrenia patients compared to healthy control groups: 0.34% of connections for ADHD, 4.71% for Yale and 0.02% for COBRE schizophrenia data were changed (see Fig. 3b). When the two schizophrenia datasets were combined, 8.13% of connections were altered in patients compared to control subjects (FDR-corrected). Note the high similarity between unthresholded RSFC patterns between the schizophrenia datasets in Figure 3a, which helps justify combining them into a single analysis.

**Fig. 3.**
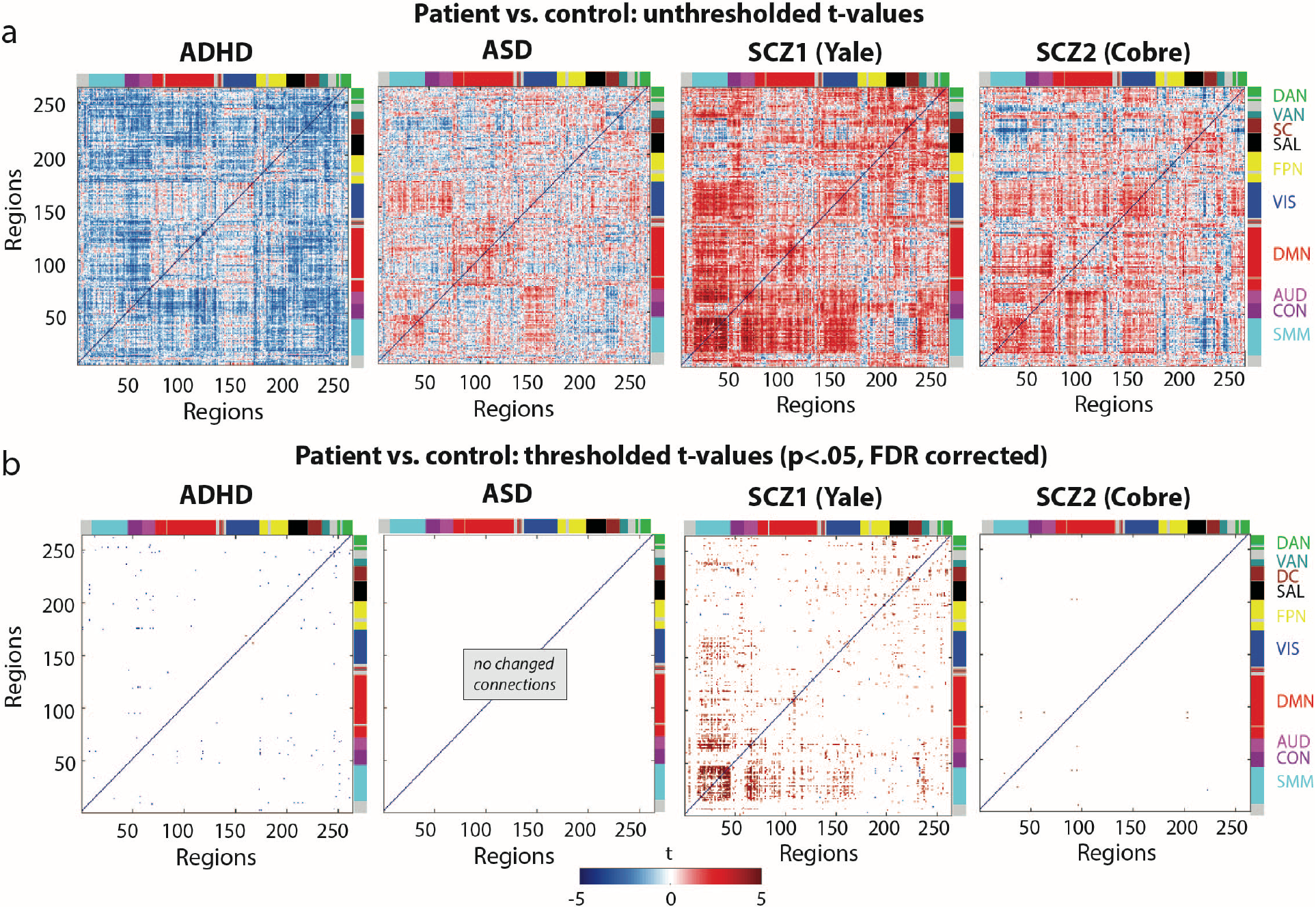
Despite high group-to-group similarity, many of the small differences between patients and healthy control subjects are consistent (as quantified by t-tests). Differences between RSFC of patients and control subjects were found for a number of connections in ADHD and schizophrenia patients (*p* < .05, FDR corrected). **a)** T-value patterns resulting from patient-control comparison of individual connections. **b)** FDR-corrected T-values (*p* < .05) resulting from the same patient-control comparison of individual connections as in panel a. Note that the smaller number of subjects for the COBRE dataset (33 patients, 45 controls) likely drove the smaller number of significant t-values relative to the Yale dataset (75 patients, 84 controls).

### 3.6 Classification based on connectome-wide similarity

Although we found functional network architecture in patients and control subjects to be highly similar, the altered connections that were found when comparing individual connections (see Figure 3) suggested connectome-wide similarity might still be informative with regard to diagnosis. We tested this idea with a classification analysis. In our classification model we used RSFC pattern similarity as a predictor variable, and tested whether an individual’s functional architecture was more similar to that of the average patient or control by using a leave-two-out approach (one patient and one healthy control subject as test set for each cross-validation fold). Note that individual-level connectome-wide similarity tended to be smaller than the group-level results, likely because group-level averaging increases signal-to-noise (from more data) and blurs across individual differences that are idiosyncratic with regard to group membership. Results from the classifications are shown in Figure 4 (chance = 50%). This analysis – based on whole-brain RSFC patterns – resulted in 58% prediction accuracy for ADHD (*p* = 0.03, 57% sensitivity, 58% specificity, N=77 per class), and 64% prediction accuracy for ASD (*p* = 0.0005, 66% sensitivity, 63% specificity, N=70 per class). For the two schizophrenia groups, we could accurately predict group membership for 75% (Yale, *p* < 0.00001, 81% sensitivity, 68% specificity, N=75 per class) and 68% (COBRE, *p* = 0.002, 79% sensitivity, 58% specificity, N=33 per class) of subjects.

**Fig. 4.**
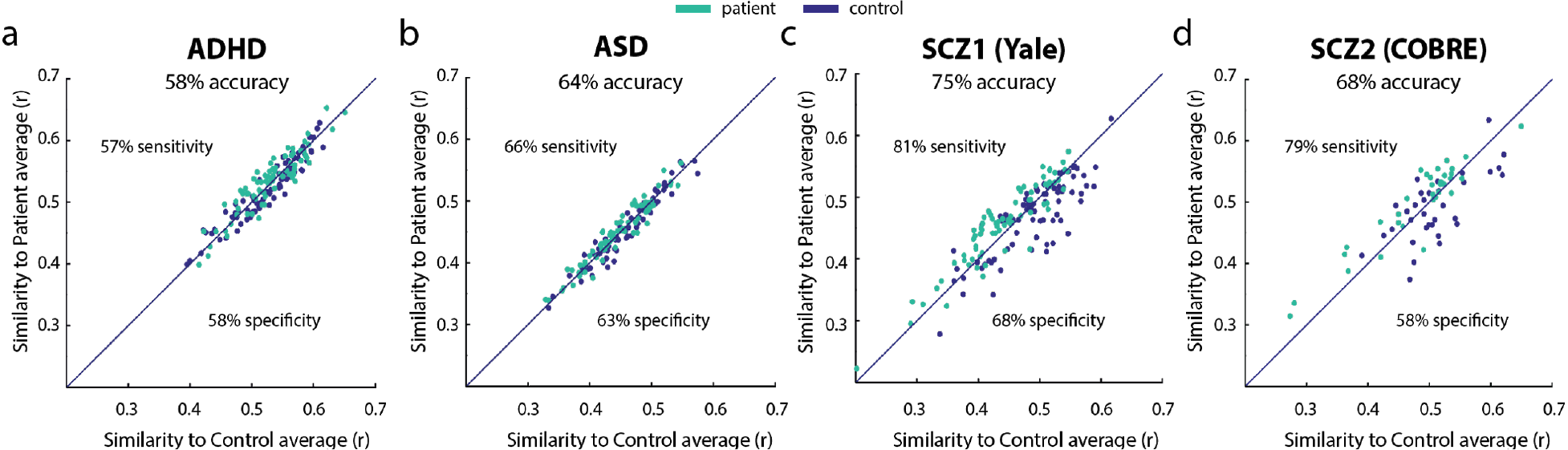
Patient-control classification based on connectome-wide resting-state functional connectivity patterns. Class labels were predicted above chance using a simple classification method in a) ADHD b) ASD, and the schizophrenia c) Yale and d) COBRE datasets. Classifying a subject involved computing that subject’s connectome-wide correlation with the patient-group average and the control-group average, with that subject being assigned to the group with the higher similarity. This demonstrated the potential informativeness of whole-brain RSFC similarity in diagnosing individuals, despite overall high similarity between patients and controls at the group level. Note that more complex classifiers would likely provide better classifications, but that straightforward inferences regarding connectome-wide correlation would not be possible with such approaches.

We next sought to test for generalization of connectome-wide RSFC patterns across datasets, assessing the ability to predict diagnoses based on cross-dataset RSFC alterations. This would suggest robustness of the RSFC alterations despite their small effect on connectome-wide similarities, and despite the data being collected at different locations (with potential scanner-specific influences on RSFC patterns). Demonstrating robust generalization, we were able to accurately predict group membership (patient vs. control) above chance across the ADHD and ASD datasets, and across the two schizophrenia datasets. For this analysis, we trained a classifier on one dataset, and tested subjects in the other dataset, and vice versa, comparing each subject’s RSFC pattern to the average control and patient FC architecture. A group label was assigned to each subject in the test dataset based on highest similarity (to either patient or to control average in the training dataset). Binomial tests showed 64% prediction accuracy (*p* < 0.000001) for ADHD and ASD datasets (N=147), with 52% sensitivity and 77% specificity. The same analysis for Yale and COBRE schizophrenia datasets (N=108) resulted in 65% prediction accuracy (*p* < 0.0001), with 68% sensitivity and 59% specificity.

## Discussion

The clinical neuroscience literature typically focuses on disease alterations in the functional connectome without putting those alterations in their overall context. We took a more comprehensive view in the present study by comparing RSFC patterns across the whole connectome, taking all connections into account at once when comparing patients and healthy controls across multiple mental disorders. By adopting such a whole-brain view of functional network architecture we aim to provide informative new constraints on psychiatric theory, emphasizing the role of small brain network changes in the negative life-altering effects of mental illness. Supporting this new perspective, we used a connectome-wide correlation analysis to demonstrate that – while being predictive of diagnosis – connectome-wide RSFC patterns are very similar across patients and controls in a range of mental disorders, and even across patient groups.

### Large overlap in network patterns across health and disease has implications for neurocognitive theories of psychiatric illness

The clinical RSFC literature to date has been so focused on showing only cross-group differences that the results illustrated in Figure 1 – showing high cross-group similarity between patients and healthy controls in ADHD, ASD, and schizophrenia – initially appear to run counter to every published paper in this area of research. Critically, however, there is technically nothing contradictory about these and prior results. These results suggest that by ignoring or understating the baseline similarity between groups, prior studies have been inadvertently presenting a biased view of psychiatric differences in brain network organization.

The observed high similarity between patients and healthy individuals has important implications for neurocognitive theories of mental illness. This is principally due to the emphasis on large-scale neurotransmitter disruptions in most psychiatric theories. For instance, current theories suggest schizophrenia may be caused by large-scale disruptions to the dopamine system (Howes and Kapur, 2009) or large-scale disruptions to the glutamate system (Moghaddam and Javitt, 2012). There are similar theories with ASD involving glutamate (Purcell et al., 2001), acetylcholine (Perry et al., 2001), and serotonin (Jr et al., 1997). With ADHD there are also similar theories involving dopamine (Kirley et al., 2002), norepinephrine (Zimmer, 2009), and serotonin (Zepf et al., 2010). All of these neurotransmitter systems are extremely widespread and critical to brain network functionality (especially glutamate), such that these theories most directly predict widespread alterations to functional brain network organization.

The current results do not rule these theories out completely, however. Rather, these results constrain these theories in important ways. It is possible that, for example, glutamatergic functionality is disrupted widely in schizophrenia, but that the disruption is minimal at each synapse. Alternatively, the disruption could substantially alter very specific functionality (such as timing of neurotransmitter binding) massively, with minimal effect on functional network organization but pervasive effects on behavior. We leave full explanations of how the present results might be compatible with observations supporting widespread neurotransmitter dysfunction in these mental disorders to future work. For now, we emphasize the need for future studies to shift their hypotheses to account for such small functional network changes.

In accounting for the small functional network changes observed here, three possibilities seem prominent: 1) Most patients actually have very similar cognition/behavior to healthy individuals in aggregate (similar routines, basic abilities like language, etc), with high brain network similarity reflecting how similar the cognition/behavior implemented by these networks actually is. The possibility is supported by the common observation of similar behavior across patients and healthy individuals (relative to random behaviors). This conclusion is also supported by the exclusion of, e.g., the most severe schizophrenia patients in fMRI studies (since they cannot comply with experiment instructions). This predicts that including severe cases of psychiatric illness would reduce the similarity between patients and controls observed here. Nonetheless, in so far as mental disorders are a matter of classification (with the most mild positive cases defining the threshold between positive and negative diagnoses), inferences based on mild cases of a disorder are likely valid. 2) Large changes in cognition/behavior arise from small changes in functional network organization. This idea is supported by the small differences in functional network organization across highly distinct cognitive/behavioral states. This high connectome-wide similarity was observed in healthy young adults across a wide variety of highly distinct tasks and resting state (Cole et al., 2014a; Krienen et al., 2014). 3) RSFC does not capture the effects of large-scale network disruption. This possibility is unlikely, given the widespread RSFC changes observed with pharmacological studies that experimentally manipulate neurotransmitter functionality (Anticevic et al., 2012; Klumpers et al., 2012; Scheidegger et al., 2012). It will nonetheless be important to test this possibility using the connectome-wide similarity approach used here. Notably, if connectome-wide similarity remains high in such studies this would also have strong implications for neurocognitive theories of mental illness, since it would suggest the brain can exhibit small functional network changes from widespread neurotransmitter functional alteration. It is also worth considering that the primary methodological issues with fMRI and RSFC is their lack of specificity (e.g., whether a signal originated from excitatory or inhibitory neurons) rather than lack of sensitivity (Logothetis, 2008; van den Heuvel and Hulshoff Pol, 2010; Ma et al., 2016; Grandjean et al., 2017), suggesting widespread functional network disruption would very likely be detected with RSFC with fMRI.

### High similarity in resting-state functional connectivity patterns between mental disorders

We also used connectome-wide correlation to compare connectivity between clinical groups, finding that they were very similar. The highest pattern similarity was found between ADHD and ASD groups (with a shared variance of 92%). This result is likely due in part to the data being collected at the same site, though even when comparing schizophrenia patients to either ADHD or ASD patients shared variance was high at 83%. Note that comparable similarity was also found for the cross-diagnostic control subjects. However, the polygenic nature of many mental disorders may also have contributed to highly similar RSFC patterns in patients. A recent study found an association between certain cross-disorder polygenic risk factors for mental disorders (ADHD, ASD, schizophrenia, bipolar disorder and major depressive disorder) and alterations in RSFC (Wang et al., 2017). Furthermore, Sprooten et al. (2016) showed that cortical regions implicated in several psychiatric disorders (based on task fMRI studies) are very similar across diagnoses, and are thus largely diagnostic-general. The present results support this possibility, extending it to include RSFC patterns shared between psychiatric disorders.

### Functionally meaningful alterations in intrinsic functional architecture of patients

The RSFC literature has clearly described alterations in FC in a variety of mental disorders, including those disorders covered in the current study. For example, schizophrenia – which is characterized by severe cognitive symptoms (Kahn and Keefe, 2013) – has been shown to involve global disruption of prefrontal cortex RSFC (Zhou et al., 2007; Cole et al., 2011; Anticevic et al., 2015). Overall, less integrated intrinsic brain networks and connections with prefrontal cortex have been found in schizophrenia patients (for a recent review of RSFC studies and links to cognition in schizophrenia, see Sheffield and Barch, 2016).

In ADHD, RSFC changes in the fronto-parietal network (FPN) have been identified and associated with cognitive symptoms like attentional control deficits, response inhibition deficits and impulsivity (Lin et al., 2015) (for a review of RSFC alterations in ADHD see Posner et al., 2014). Further supporting the involvement of FPN RSFC in ADHD, one study was able to predict individual IQ scores (a reduction in IQ is strongly associated with ADHD) based on FPN connectivity measures in children and adolescents with ADHD (Park et al., 2016). Dysfunctional connectivity of cognitive control networks such as FPN to the default-mode network have also been implicated in ADHD (Castellanos et al., 2008; Sun et al., 2012; Hoekzema et al., 2014). The extent to which these changes represent fundamental alterations to brain system organization remains unclear, however.

Patients with ASD show cognitive impairments in several domains such as social cognition, language, attention, executive function and working memory (Baron-Cohen et al., 1985; Rogers and Pennington, 1991; Charman et al., 2011). RSFC studies have indicated a variety of functional network changes associated with ASD, with these changes including both over-connectivity and under-connectivity in areas associated with the cognitive impairments seen in ASD (for a review see Hull et al., 2017).

These RSFC alterations identified by previous studies might seem to contradict our results indicating little differences between patients and healthy controls. We therefore performed a series of analyses to test whether the overall theme of prior results (that there are reliable RSFC differences between patients and control subjects) could be replicated in the data used here. The results indicated that the whole-brain FC pattern can be largely similar across patients and control subjects while there simultaneously being small but reliable RSFC alterations consistent with the existing literature.

When testing all individual connections, alterations survived correction for multiple comparisons (with the exception of ASD), and thus showed the existence of deviant RSFC architecture in the current study. Interestingly, t-statistic pattern correlations (Fig. 3A, connectome-wide patterns of unthresholded t-test results) between mental disorders were significantly above chance, indicating there is a shared component in the deviant functional connections between disorders. Correlations were largest between ADHD and ASD, and between the Yale and Cobre schizophrenia datasets, which is likely a combined effect of data collection site and diagnosis (note that there is high comorbidity between ADHD and ASD (Leitner, 2014)).

Further reconciling the present results with the existing clinical neuroscience literature, we used machine learning to show that the small connectome-wide RSFC differences between patients and healthy controls were nonetheless predictive of clinical status. Specifically, we found above-chance classifications of patients vs. healthy controls in all three clinical groups, based on connectome-wide correlation of RSFC values. This resulted in 58% clinical status prediction accuracy for ADHD, 64% for ASD, 75% for the Yale schizophrenia dataset, and 68% for the COBRE schizophrenia dataset. Notably, the connectome-wide correlation differences used to make these predictions were quite small (see Figure 4), consistent with the small cross-group connectome-wide correlation differences identified in the main results. Together, these results further reconcile the current results with the rest of the clinical RSFC literature, demonstrating the utility of RSFC pattern changes despite the small overall connectome-wide differences associated with the psychiatric diseases investigated here.

## Acknowledgements

We acknowledge support by the US National Institutes of Health under awards K99-R00 MH096801, R01 AG055556, and R01 MH109520. The content is solely the responsibility of the authors and does not necessarily represent the official views of any of the funding agencies.

